# Examining and fine-tuning the selection of glycan compositions with GlyConnect Compozitor

**DOI:** 10.1101/2020.06.03.131979

**Authors:** Thibault Robin, Julien Mariethoz, Frédérique Lisacek

## Abstract

A key point in achieving accurate intact glycopeptide identification is the definition of the glycan composition file that is used to match experimental with theoretical masses by a glycoproteomics search engine. At present, these files are mainly built from searching the literature and/or querying data sources focused on posttranslational modifications. Most glycoproteomics search engines include a default composition file that is readily used when processing mass spectrometry data. We introduce here a glycan composition visualising and comparative tool associated with the GlyConnect database and called GlyConnect Compozitor. It has web interface through which the database can be queried to bring out contextual information relative to a set of glycan compositions. The tool takes advantage of compositions being related to one another through shared monosaccharide counts and outputs interactive graphs summarising information searched in the database. These results provide a guide for selecting or deselecting compositions in a file in order to reflect the context of a study as closely as possible. As part of the tool collection of the Glycomics@ExPASy initiative, Compozitor is hosted at https://glyconnect.expasy.org/compozitor/ where it can be run as a web application. It is also directly accessible from the GlyConnect database.

## Introduction

One of the most interesting current challenges in proteomics research is estimating the extent of protein diversity as recently summarised in (1). To encompass all molecular forms of a protein following genetic variation, alternative splicing and posttranslational processing, the term “proteoform” was recently proposed (2) and rapidly adopted by the community. Yet, the array of posttranslational modifications is not fully characterised and the particular case of glycosylation raises many technical issues that glycoproteomics has started to address on a large-scale basis only a few years ago. Recent examples include cancer (3) (4) or mouse brain (5) N-glycome profiling studies. These studies rely on proteomics software adapted to identifying glycopeptides such as Mascot (6) and ProteinProspector (7) or glycoproteomics dedicated software developed in recent years as reviewed in (8).

A key point in achieving accurate glycopeptide identification is the selection of a glycan composition file that will be used to match experimental with theoretical masses as is usually implemented in mass spectrometry (MS) search engines. The definition of this composition file differs in the range of software commonly used. It is generally selected to cover as many monosaccharide combinations as possible and is seldom customised to reflect the expectable constraints imposed on glycan expression in a specific tissue or organism. For example, in the very popular Byonic search engine (9), the default list of N-glycan compositions is set to 309 items and usage reported in publications more often than not shows a selection of default values when human samples are processed. Nonetheless, the list can be customised and for example, the mouse brain study (5) describes the selection of compositions extracted from the literature with relevant criteria (e.g., relevant species and tissue). At present, searching the literature and/or querying other data sources such as Unimod (http://www.unimod.org) appear to be the most frequent approach to customising a composition file.

In 2017, the HUPO Human Glycoproteomics Initiative launched an interlaboratory study to assess the performance of glycoproteomics software for automated intact N- and O-glycopeptide identification from high resolution MS/MS data. Benchmark datasets were provided to participants split as developers (write software) or users (use software tool selected from the range of existing ones). Data was analysed and the results sent back to the challenge organisers for evaluation. This revealed widespread variations in the definition of composition files and the subsequent variations in the quality and extent of glycopeptide identification.

We describe here a web-based tool destined to assist glycoproteomics software users in selecting appropriate N- or O-linked glycan compositions with respect to sample specifications encompassing species, tissue or cell line type and disease. This tool named *Compozitor*, relies on the data collected in the GlyConnect resource (10), which includes glycomics and glycoproteomics data. In fact, Compozitor reveals global glycomic information associated with a species, a glycoprotein, a cell or a tissue that cannot be captured when reading through the corresponding GlyConnect entries. In particular, a glycome is often provided as a list of glycan structures or a list of compositions or a mix thereof, as if the items were independent when they obviously are not. The interface of GlyConnect described in (10) was a first attempt to link and visualise glycomic and proteomic data, e.g., addressing the question of which glycan(s) is/are attached to which protein(s). Navigation in the database did not support the comparative investigation of structural data dependencies, e.g., addressing the question of detecting glycan compositional trends within and similarities across protein(s) or tissues.

As part of the tool collection of the Glycomics@ExPASy initiative (11), Compozitor is hosted on the ExPASy server (12) of the SIB Swiss Institute of Bioinformatics at https://glyconnect.expasy.org/compozitor/ where it can be run as a web application. It is also referenced in GlyConnect glycoprotein, source, reference and disease pages to offer a new view on the data. It appears in the cross-reference section of these pages. The present article describes the tool and demonstrates its use through a series of use cases arising from the exploration of the GlyConnect database content. It also tackles the comparison of composition files used in several intact glycopeptide search engines and suggests options for rationalising a selection.

## Material and Methods

### GlyConnect glycosylation data

The GlyConnect database stores curated data on glycosylation, glycans and glycoproteins extracted from literature. It was built upon the wealth of information contained in the GlycoSuiteDB database (13). GlyConnect was then enriched with data and annotations provided by Nicki Packer’s group through collaborative work (10) (11) and by the recent integration of selected published work on high throughput mass spectrometry experiments using a range of identification software in various applications. The May 2020 release of GlyConnect contains 246 species, 2663 proteins, 5675 glycosites, 1041 compositions, 3607 defined and 451 ambiguously defined structures. There are twice as many N- than O-linked recorded in general and as a reflection of the current bias in the recent literature, 53% of site-specific data corresponds to human N-linked glycans. Note that 6% of the reported glycosylation is relative to released glycans.

The data is provided to users through a web application. Two types of representation are available. The application is either serving HTML web pages for human readability or JSON (JavaScript Object Notation) with a RESTful API (Application Programming Interface) for software applications. The latter is used by GlyConnect Compozitor to populate the menu options and return the results of the composition search query. The RESTfull API can also be used to convert different composition notation formats or return GlyConnect or GlyTouCan identifiers as well as monoisotopic masses.

### Composition notation

There is no single way of representing glycan compositions and Compozitor currently implements three of the most common notations, especially in mammalian studies. The residue set is composed of monosaccharides and substituents (see Table 1). In contrast to resolved glycan structures, the number of monosaccharides in compositions is reduced to a smaller set of residues that have identical molecular masses (e.g.180 Da for one hexose that is valid for galactose and glucose).

**Table 1:**
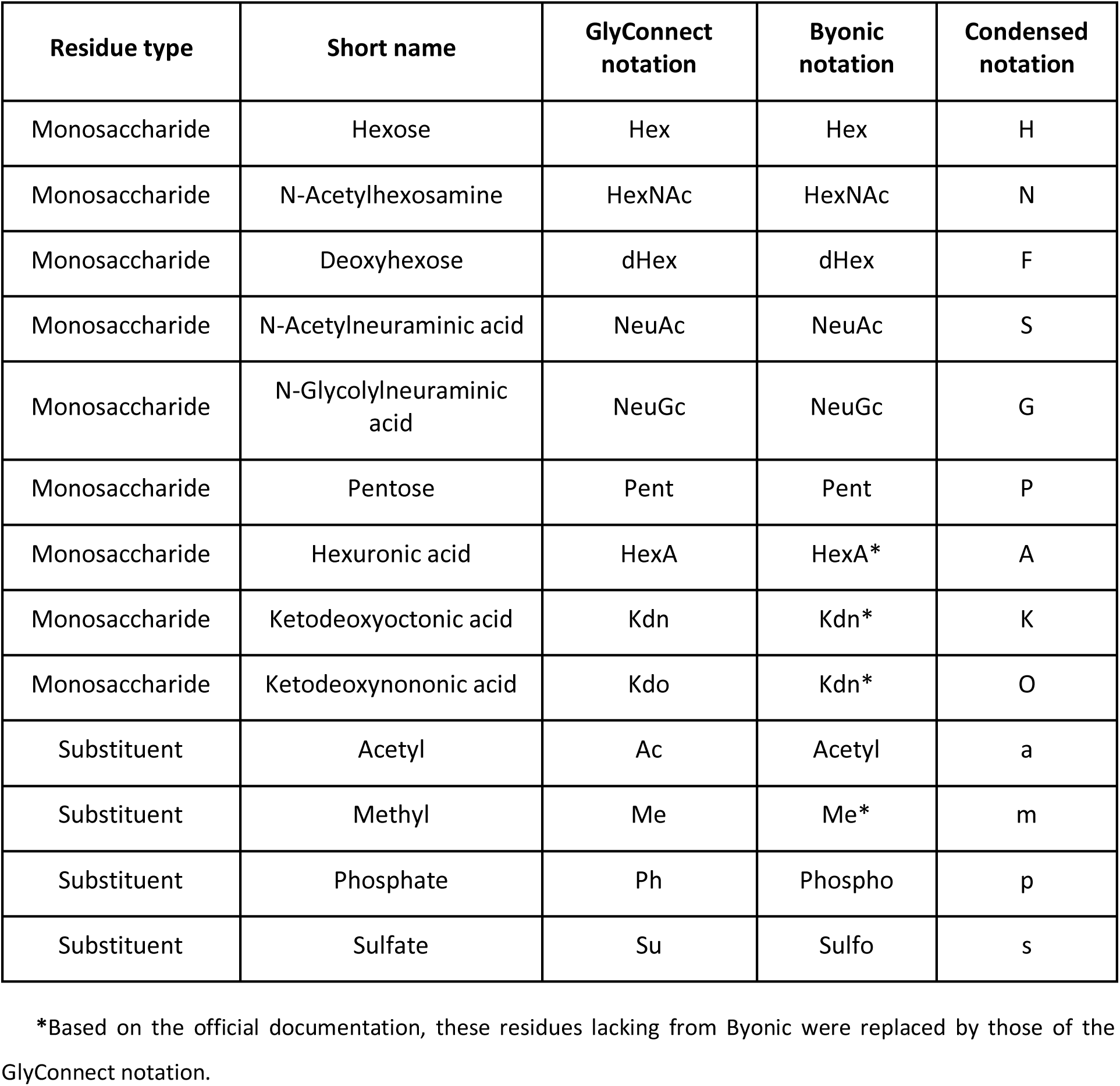
Common glycan composition residues and corresponding abbreviations

| Residue type | Short name | GlyConnect notation | Byonic notation | Condensed notation |
| --- | --- | --- | --- | --- |
| Monosaccharide | Hexose | Hex | Hex | H |
| Monosaccharide | N-Acetylhexosamine | HexNAc | HexNAc | N |
| Monosaccharide | Deoxyhexose | dHex | dHex | F |
| Monosaccharide | N-Acetylneuraminic acid | NeuAc | NeuAc | S |
| Monosaccharide | N-Glycolylneuraminic acid | NeuGc | NeuGc | G |
| Monosaccharide | Pentose | Pent | Pent | P |
| Monosaccharide | Hexuronic acid | HexA | HexA* | A |
| Monosaccharide | Ketodeoxyoctonic acid | Kdn | Kdn* | K |
| Monosaccharide | Ketodeoxynononic acid | Kdo | Kdn* | O |
| Substituent | Acetyl | Ac | Acetyl | a |
| Substituent | Methyl | Me | Me* | m |
| Substituent | Phosphate | Ph | Phospho | p |
| Substituent | Sulfate | Su | Sulfo | s |

\*Based on the official documentation, these residues lacking from Byonic were replaced by those of the GlyConnect notation.

The GlyConnect notation is inherited from one of the oldest software tools managing compositional data known as GlycoMod (14). It uses an abbreviated code for residues and a semicolon as a separator between this code and the corresponding number of monosaccharides. Then each such pair is separated from the next by a space. For example, the N-glycan core made of two N-Acetylhexosamine and three hexoses is represented as: Hex:3 HexNAc:2.

The Byonic notation is included to ease export from the interface to a file directly importable in the analysis software for customising compositions. It uses an abbreviated code for residues very similar to the GlyConnect notation, but the corresponding number of residues for this code is between brackets and no space separates code-number pairs. The N-glycan core is represented as: Hex(3)HexNAc(2).

Finally, the condensed format uses a one-letter code immediately followed by the corresponding number of residues and does not use separators for code-number pairs. The N-glycan core is represented as: H3N2. In contrast to the two other formats, the condensed notation is case-sensitive with the monosaccharide using uppercase letters and substituents using lowercase letters. As a consequence, some residues may share the same letter with a different case (e.g., upper S for N-Acetylneuraminic acid and lower s for sulfate).

The order in which the residue is reported is consistent. It starts with the hexoses, followed by N-Acetylhexosamine, deoxyhexoses, sialic acids and substituents in all notations. Of note, this order is a tacit consensus with no biological relevance attached to it.

In the rest of this article we shall refer to the GlyConnect, Byonic and condensed notations to specify which format is being used in which context.

### Search criteria

Search criteria correspond to expected sources of variation in glycan expression such as species, tissue or disease, and they also reflect the information stored in the GlyConnect database. The interface of Compozitor offers six tabs labelled with search criteria, namely: protein, source, cell line, disease, custom and advanced.

In the *protein, source* and *disease* tabs, a “species” drop-down list can first be used to select the relevant species. This selection step is skipped in the case of cell lines, as they represent only a small number of entries. Once the species is chosen, the available options for the corresponding biological entities are proposed in a second drop-down list. In the case of the source tab, three distinct drop-down lists are provided corresponding to the three biological source categories available in GlyConnect: tissue, cell type and cell component. Of note, several sources can be selected at the same time when available. In a third step, a “glycan type” drop-down list enables to filter the retrieved glycan compositions based on the type of linkage (i.e., N-linked, O-linked or C-linked). In the protein tab specifically, an additional “sites” drop-down list is instantiated each time a glycan type is selected, displaying all known glycosylation sites of the corresponding protein. Compositions recorded on these sites are all included by default yet, any site can be checked or unchecked by the user. These four tabs all depend on the “glycosylations” public route of the GlyConnect RESTful API to retrieve the corresponding glycosylation information. Direct access to this route is provided via the *advanced* tab, where specific queries can be input in the form of URL parameters as detailed in the corresponding API help page:https://glyconnect.expasy.org/api/docs.

To expand the usage of the tool beyond the content of the GlyConnect database, a *custom* tab was implemented. It allows inputting a composition list in either of the three supported formats by copy-pasting in the appropriate text field or uploading a text file. These custom compositions are mapped to those contained in GlyConnect, in order to provide basic information for each match, such as GlyConnect and GlyTouCan identifiers, molecular masses and known glycan structures. These conversion and mapping services can also be used externally as described in the API help page.

In either of the selected tab, once the different search criteria have been chosen, the query is activated by clicking on the “Add to selection” button. This selection is visually summarised as a combination of search criteria (i.e., the species, biological entity, glycan type and glycosylation sites for proteins), followed by the corresponding number of glycan compositions recorded in the GlyConnect database. Each biological entity is cross-linked via an accession number to its matching reference database/ontology as detailed in Supplemental Table 1. A given selection can be discarded at any time by clicking on its cross icon.

In the current version of Compozitor, up to three search results can be simultaneously active. In other words, the respective known glycomes of multiple combinations can be compared, such as, three proteins or two proteins and a tissue, etc. Each search result is assigned a label in the form of a block letter. The chronological order of successive queries is matched with the alphabetical order of block letter labels (A represents the first, B the second and C the third set of results). The “Compute graph” button triggers the graphic display of glycan compositions selected by queries and enables further investigation of the results. Alternatively, users only interested in downloading composition lists can use the “Export selection” option.

### Graphical result display

Frequently enough, a glycan may be fully contained in one or more other compositions. To account for this dependency, the search results are displayed as a directed graph in which each node is a composition and two nodes are connected if they differ by one residue (as listed in Table 1). The graph generation is powered by D3.js (https://d3js.org), a flexible JavaScript library providing data visualization for web applications. The layout of the graph is based on the force-directed algorithm implemented in D3.js. When queries yield a large number of unique compositions, all computations are performed in background threads using *web workers*, i.e., protocols for web pages to execute tasks in the background. This prevents the user interface from becoming unresponsive when a graph contains hundreds of nodes.

#### Nodes

The graph nodes represent the unique set of glycan compositions contained in one or several query results. A single set of results is labelled A and shown with blue nodes. In the case of multiple queries, each set of results is assigned a distinct colour (blue A, red B and green C). When a composition is shared between several sets of results another colour scheme is used (magenta AB, yellow AC, cyan BC and black ABC). A legend located on the upper left side of the display keeps track of the search criteria, as well as the colour and label codes. The occurrence count of each specific colour node in the graph is shown in the legend as the number that labels each coloured node.

In the graph, each node is identifiable with a glycan composition in the condensed notation. Other features reflect information stored in the GlyConnect database. Each node is labelled inside with the number of glycan structures matching the corresponding composition in the context of the search criteria. Node size varies based on the number of publications in which the composition was detected. Mousing over a node label prompts a popup window containing finer details of the composition: cross-links to GlyConnect and GlyTouCan (15), the monoisotopic mass in Daltons, and related glycan structure cartoons in the Symbol Nomenclature for Glycans (SNFG) format (16). A small window offering a “zoom on” opens a drop-down list of all nodes of the graph categorised as *root, leaf, unconnected* and *other*. It is located on the top right of the main display to facilitate searching for a particular composition in the graph especially when search criteria generate a large and crowded graph in which specific nodes may be difficult to find. Once a given composition is selected, the graph view centers on the corresponding node.

One of the key features of Compozitor lies in the addition of grey “virtual nodes” to increase the connectivity of a graph. These nodes are computed by performing the systematic pairwise comparison of all compositions in a graph. If two compositions differ from exactly two residues, the two corresponding nodes are tentatively connected through an intermediary node that is only one residue away from each. Then, this tentative node is added only if it meets two conditions:

∘ It does not already exist in the graph
∘ None of its children has a non-virtual parent node (i.e., an alternative path is possible in the graph to connect two nodes differing from exactly two residues).

The second selective rule is implemented so as to avoid overcrowding the graph with unnecessary nodes.

#### Paths

The edges of the graph are represented as grey unidirectional arrows showing the addition of a residue. Similarly to nodes, each edge is labelled with the letter of the condensed notation corresponding to the added residue. To evoke the SNFG colouring scheme, the addition of one fucose is displayed in SNFG red and that of one sialic acid in SNFG purple. Although less strictly defined, SNFG blue was assigned to the addition of one N-Acetylhexosamine and SNFG green to that of one hexose.

In an attempt to estimate the connectivity of the graph and the consistency of relatedness between compositions, the reachability of each node is also shown when mousing over it. Entering paths are highlighted in cyan while exiting paths in orange. Each node can be moved interactively by dragging it in the desired direction. It is also possible to follow paths associated with the addition of a specific residue. Mousing over a path labelled with the monosaccharide that is added will trigger the highlight of all paths labelled with that particular monosaccharide addition. Finally, zoom buttons are available at the bottom of the page for magnifying or reducing the graph.

#### Glycan properties

Glycans of the GlyConnect database are associated with general properties commonly used to qualify or categorise them. In the current version of Compozitor, three of these (*neutral, fucosylated, sialylated*) qualify compositions irrespective of their type and are complemented with *oligomannose* as N-glycan specific and *sulphated* as O-glycan specific. Then, five properties are assigned to either type of glycan compositions according to rules defined upon the advice of glycobiologists and detailed in Table 2.

**Table 2:** Reported glycan properties by linkage type

| Glycan type | Properties | Rules |
| --- | --- | --- |
| N-linked/O-linked | *neutral* | does not contain any sialic acid (N-Acetylneuraminic acid (NeuAc) or N-Glycolylneuraminic acid (NeuGc)) or sulphate (Su) |
| N-linked/O-linked | *fucosylated* | contains at least one deoxyhexose (dHex) |
| N-linked/O-linked | *sialylated* | contains at least one sialic acid (N-Acetylneuraminic acid (NeuAc) or N-Glycolylneuraminic acid (NeuGc)) |
| N-linked/O-linked | *fuco-sialylated* | contains at least one deoxyhexose (dHex) AND at least one sialic acid (N-Acetylneuraminic acid (NeuAc) or N-Glycolylneuraminic acid (NeuGc)) |
| N-linked | *oligomannose* | contains at least 5 hexoses (Hex), no more than two N-Acetyl hexosamine (HexNAc) and no more than one deoxyhexose (dHex) |
| O-linked | *sulphated* | contains at least one sulphate (Su) |

Glycan properties are directly inferred from the presence/absence of specific monosaccharides in compositions except. the oligomannose property that is deduced from monosaccharide counts and as such, is not as reliably assigned as the others. Note that *fucosylated* is assigned upon the presence of a deoxyhexose, but deoxyhexoses may correspond to monosaccharides other than fucose (e.g., quinovose or rhamnose) in some non-mammal species. It is consequently correct for most entries in the GlyConnect database, but can be erroneous in a few specific cases.

A global view of the compositions contained in the graph is captured in an interactive bar plot of the five-property counts, displayed next to the graph on the bottom right. Mousing over any bar of the bar plot highlights in orange all nodes contributing to the corresponding frequency.

#### Custom datasets

As mentioned earlier, the user is given the option of inputting custom compositions. In that case, launching the search involves a systematic comparison with the GlyConnect database content irrespective of any selection criterion other than matching the input compositions. Each item of the input list is searched in the database composition entries and when a match is found, the corresponding information is kept to be mapped on the graph. In the end, a graph is generated with nodes that are exactly those of the input list. Their respective size reflects the number of recorded publications supporting the corresponding composition and their label is the number of solved structures stored in the database for the corresponding composition. In this way, the graph provides a rough estimate of how realistic a composition set may be.

### Export

The glycan compositions of each selected query can be exported as a text file in all supported formats. As mentioned earlier export can be launched before the generation of the graph to create composition lists, but when it is launched after displaying the graph, all compositions or composition subsets can be selected. More precisely, compositions that are common to several sets of results or those that correspond to virtual nodes can be singled out, therefore saved or discarded. In all cases, basic metadata information is added in the form of a header at the beginning of the exported files. It describes the tool version and GlyConnect release, date of generation, composition format, and selected queries. A shortcut copy icon also allows a quick copy-paste in the clipboard. Additionally, the graph may be exported as a vectorial image in the SVG format.

## Results and Discussion

Most of the results presented in this section were obtained with human N-glycomes simply because the vast majority of current glycoproteomics publications have this focus. Consequently, sampling problems tend to be minimised in this case.

### Visualisation and interpretation of a glycome graph

The description of the tool in the Material and Method section emphasises the array of information that is displayed in the Compozitor interface. In particular, the graph representation of the glycome of a protein, a tissue or a cell line is more amenable to capturing potential compositional biases.

The bar plot of N-glycan overall properties is the first summary view. In absence of a reference for an expected distribution depending on species or tissue, it is at best informative on its own and mostly useful in the comparative mode.

Even though the output graphs tend to simplify the reality of the underlying enzymatic network at work in synthesising glycans, this representation may reveal (dis)continuous paths and provide clues regarding the likelihood of a structure in a given context. Common sense would suggest that if a composition occurs in the graph, it will stem from compositions that have one less monosaccharide. This is precisely the information brought by the graph. In the end, connectivity reflects the accessibility of a node via other nodes and emphasises the consistency of synthetic reactions. From a formal point of view, node reachability provides a classical estimate of the graph properties. For example, some GlyConnect protein records contain associated compositions that produce scattered and poorly connected nodes while others generate a fully connected graph. The introduction of virtual nodes plays an important role in bridging the gaps. Examples illustrating these variations are shown in Figure 1. Figures 1A-C show the N-glycome graphs of several well characterised human glycoproteins: alpha-fetoprotein (P02771), coagulation factor XI (P03951) and interferon gamma (P01579). These examples highlight both the variety of N-glycomes as confirmed by the variability of property bar plots and their respective consistency. Note that these examples mainly emphasise that with a similar size (about 20 nodes) the introduction of virtual nodes has distinct effects. No virtual node can enhance the tightly connected graph of alpha-fetoprotein (H4N2F1 remains isolated because this composition is too distant from all others). Only two virtual nodes are needed to join two clusters of coagulation factor XI, while four are needed to connect the three clusters of interferon gamma.

**Figure 1:**
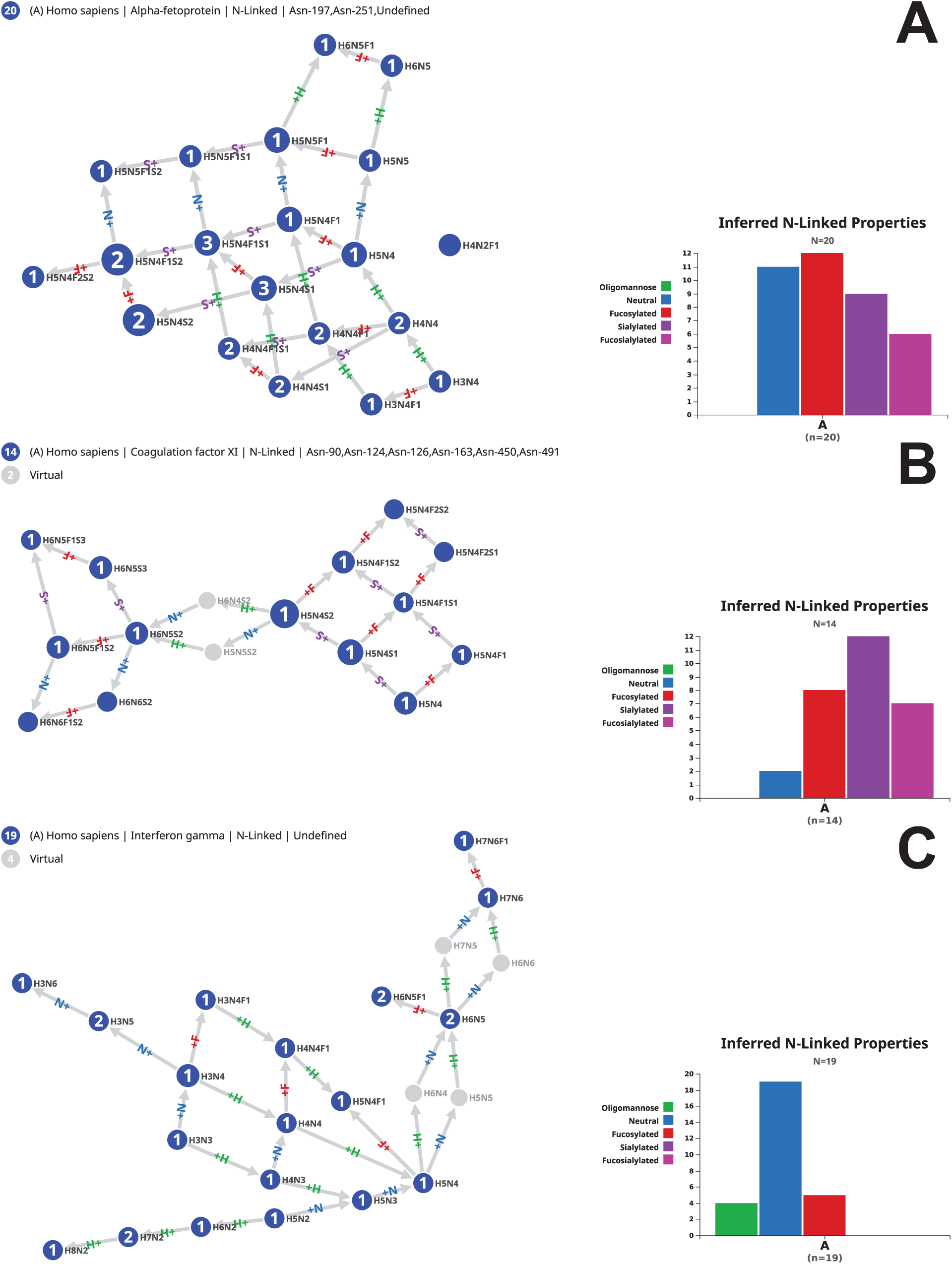
Examples of protein N-glycomes and connecting role of virtual nodes. (A) N-glycome of alpha-fetoprotein (P02771) where virtual nodes are not needed to fully connect corresponding compositions, (B) N-glycome of coagulation factor XI (P03951) where only two virtual nodes are needed to fully connect corresponding compositions, (C) N-glycome of interferon gamma (P01579) where four virtual nodes are needed to fully connect corresponding compositions. In each case the bar plot of glycan properties is distinctive.

The glycome size does not correlate with an increase in virtual nodes. Supplemental Figures S1A-B show the larger N-glycomes of human thrombospondin-1 (P07996) and non-recombinant erythropoietin (P01588). The 83 compositions of thrombospondin-1 form an almost fully connected graph leaving out two nodes: H3N2 and H12N2. The virtual node completion integrates the former in the graph through connections to initially missing H3N3 and H4N2. However, H12N2 is three monosaccharides away from H9N2, the terminal node of the path adding hexoses to H3N2. The 58 compositions of erythropoietin form scattered clusters that require 16 virtual nodes to transform the graph into an almost complete one that only leaves out a sulphated as well as an acetylated composition.

Computed graphs are interactive and can be scrutinised to detect the potential pivotal role of some nodes. A composition can be central in one graph and a root or a terminal leaf in another. This is illustrated in Supplemental Figure S2AB where H6N5F1S2 obviously plays a different role in the respective N-glycome graphs of human (A) erythropoietin (P01588) and (B) decorin (P07585). The removal of H6N5F1S2 in (A) would break six connections in the erythropoietin graph thereby disrupting a path between H7N6F1S2 and H6N5F1S1 (virtual nodes H7N5F1S2 and H6N6F1S2 would be unjustified) and would create an isolated cluster with only H6N5F1S3 and H6N5F1S3a1. In contrast, discarding H6N5F1S2 in (B) is of minimal consequence since all paths are preserved and only H6N5F1S3 is left as an isolated node.

Likewise, a node may be virtual in a graph and steadily mapped in another. We interpret the oscillation of a node between virtual and real in similar contexts as indicative of the relevance of its role. In some cases, a missing composition may point at missing data and suggest a possible check of experimental data where the composition may have been below threshold in processed data results. An example is given in Supplemental Figure S3AB. H4N4F2 is virtual in (A) the extracellular matrix (ECM) protein decorin (P07585) and real in (B) thrombospondin-1 (P07996) also an ECM protein. Nonetheless, the two nodes play a comparable role with a similar connectivity in the graph. Ten cyan (outward) and 12 orange (inward) links stem from H4N4F2 in the thrombospondin-1 graph while in the decorin graph, H4N4F2 gives rise to nine cyan and 11 orange links. Furthermore, when virtual nodes are not included, then H4N4F2 is not included in the N-glycome graph of human decorin and H4N4F1 and H4N4F3 are remote as visible in Supplemental Figure S3C. No path joins them and they are therefore unreachable from one another. However, as shown in Supplemental Figure S3D, when H4N4F2 is introduced as one of the seven virtual nodes, H4N4F1 and H4N4F3 are logically connected through the H4N4F2 virtual node. In the end, not only is H4N4F2 as virtual in one glycome, comparable to its real counterpart in another glycome but when included as virtual, it connects nodes that should be mutually reachable in the graph. This suggests that H4N4F2 is likely to be real in the decorin glycome and may just have been below threshold in glycoproteomics results.

Furthermore, path highlighting reveals the contribution of a node to the graph. For example, some nodes bridge two parts of the graph as H5N4F1 occurring in the graph of the N-glycome of decorin shown again in Supplemental Figure S4A. Three paths end on H5N4F1 and four start from it. This highlights that when multiple routes go through one node, the impact of removing that node may substantially alter pathways. This information is often key to appraising the role of the corresponding composition. In contrast, H5N4F2S1 in the same graph shown in Supplemental Figure S4B is manifestly terminal. All paths end on that node and do not extend any further. In that case, the role of the node is more difficult to interpret yet it highlights the reality of node categories mentioned earlier, namely, central, root or terminal.

Note that in many of the figures, we have used the flexible interface to move a few nodes around to unpack dense regions and improve readability.

### Comparison of entities

#### Protein – Protein

The GlyConnect database contains ∼2600 protein records and comparing their respective glycomes can be informative. For example, several protein entries describe the glycosylation of integrins. In particular, as reported in the literature and transcribed in the corresponding UniProtKB/Swiss-Prot entries integrin alpha 5 (P08648) and integrin beta 1 (P05556) form a complex (ITGA5:ITGB1) that acts as a receptor for fibronectin, fibrinogen and fibrillin-1 among other recorded functions. The glycosylation of the complex was studied (17) and the results are reported in the corresponding GlyConnect entry (ID: 283). High throughput glycoproteomics experiments also report the identification of intact glycopeptides of integrin alpha 5 (ID:1407) and integrin beta 1 (ID: 1414) separately. We compared the N-glycomes of the three GlyConnect entries to estimate the overlap between these independent experiments. The results are shown in Figure 2. The bar plot in Figure 2A emphasises the distinct profiles of each N-glycome in respectively integrin alpha 5 (21 compositions mostly sialylated), integrin alpha 5/beta 1 (23 compositions) and integrin beta 1 (43 compositions mostly neutral and fucosylated). The legend of the graph (top left) indicates poor overlap which is to be expected between integrin alpha 5 and integrin beta 1 but less between the complex and each participant. This trend is confirmed by the high number of virtual nodes required to connect the clusters visible in Figure 2B into a larger graph shown in Figure 2C. The introduction of 24 grey/virtual nodes in Figure 2C increases the connectivity by 40%. As an example, the reachability of H6N5 is highlighted in both Figure 2B and C. In Figure 2B, H6N5 is reaching out to 14 nodes (orange paths) but cannot be reached from any other node. In Figure 2C, H6N5 can be reached in several steps from H5N2 (cyan paths) and links to 26 non-virtual nodes (orange paths). In all cases, the peripheral role of green nodes of integrin beta 1 is noticeable. In contrast, compositions shared between datasets play a central role.

**Figure 2:**
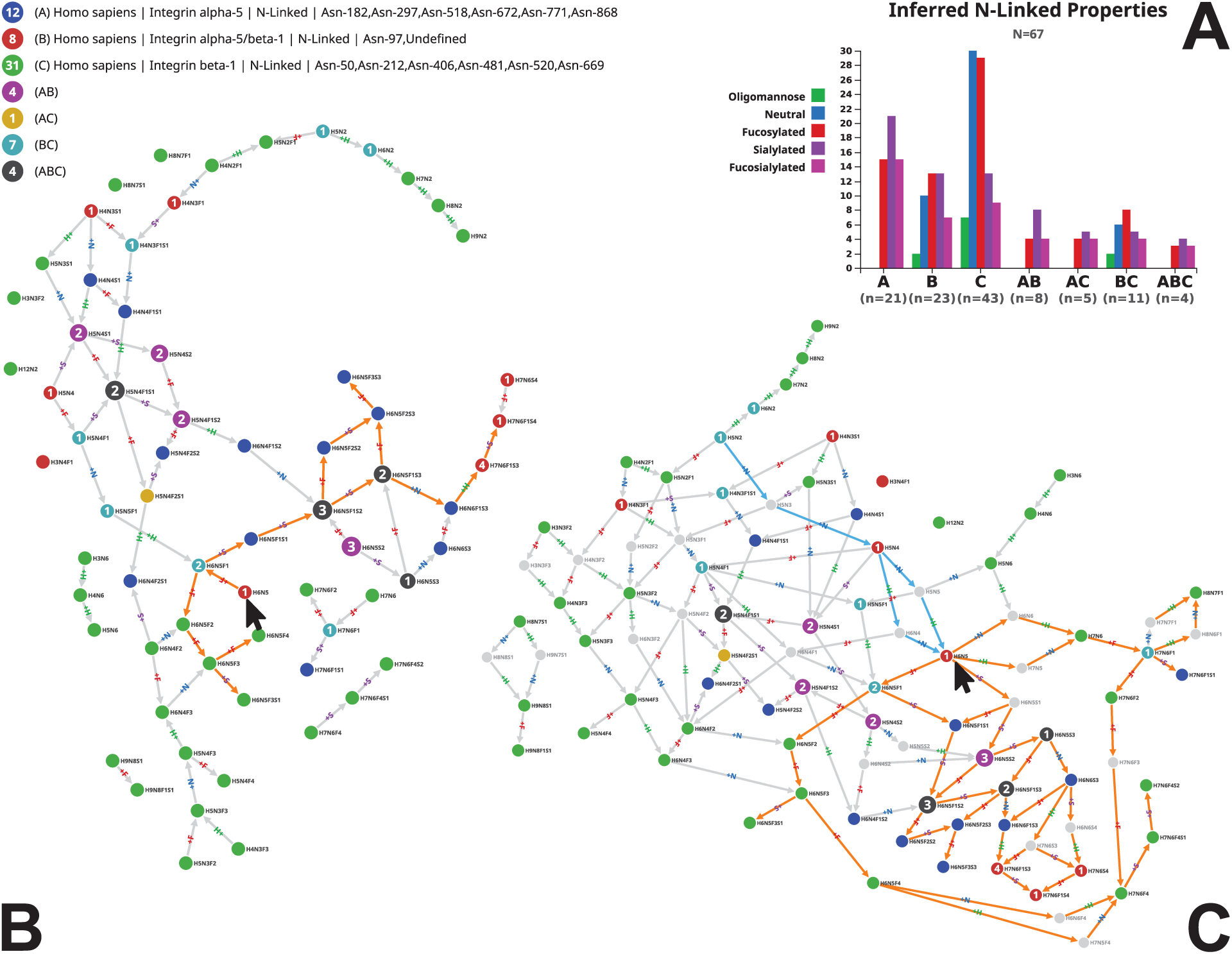
N-glycomes of two individual and complexed integrin chains. GlyConnect protein entry ID 283 describes the glycosylation of the alpha 5/beta 1 integrin complex while protein entries ID 1407 and ID 1414 separately report the individual glycosylation of integrin alpha 5 and integrin beta 1 identified as intact glycopeptides. The N-glycome of each of the three entries were compared. (A) The N-glycome properties of each protein entry are plotted and display different distributions. (B) The resulting graph with no virtual nodes displays many clusters and limited accessibility of nodes, highlighted with the status of node H6N5 (arrow) only linking out to 14 nodes via orange paths. (C) The connectivity of the resulting graph with 24 virtual nodes is enhanced as highlighted with the status of H6N5 (arrow) accessible from two real and two virtual nodes (cyan paths) and reaching out to 26 nodes via orange paths.

Finer grain information of glycosylation can be visualised through selecting one site at a time in the interface. Supplemental Figure S5 shows the comparison of conserved Asn-72 in human alpha-1 acid glycoprotein 1 (P02763) and alpha-1 acid glycoprotein 2 (P19652) that are 89.5% identical in amino acid sequence. The graph shows that the 19 compositions reported on Asn-72 of alpha-1 acid glycoprotein 2 are included in the 26 reported on Asn-72 of alpha-1 acid glycoprotein 1. The seven compositions unique to alpha-1 acid glycoprotein 1 are either neutral or sialylated. The latter tend to be terminal and the former add consistency to the graph by providing a new root and multiple connecting paths via virtual nodes.

Other combinations, such as mapping the glycome of protein with that of a specific tissue or a specific disease, can be explored. The integration of Compozitor in GlyConnect reference pages provides the display of results in each published article stored in the database. Interestingly, the corresponding graphs are often consistent and do not require the addition of many virtual nodes (data not shown).

### Consistency of well-characterised glycomes

We have used Compozitor to estimate both the extent and the consistency of some of GlyConnect glycomes. We illustrate this approach with two examples representative of glycome mapping, i.e., the N-glycome of the CHO (Chinese Hamster Ovary) cell lines and that of the human immunoglobulin gamma (IgG).

#### N-glycome of CHO cell lines

The N-glycome of the generic CHO (CVCL_0213) cell line included in GlyConnect is made of 79 N-glycan compositions summarising the information extracted from 27 publications. This data is visualised in Compozitor by querying “CHO” in the cell line tab. In the resulting graph shown without virtual nodes in Figure 3A maps the 79 compositions in three clusters and a few isolated nodes. Introducing virtual nodes generates an almost fully connected graph except for the N1F1 disaccharide that remains several monosaccharides away from other compositions/structures, as shown in Figure 3B. Sixteen virtual nodes are necessary to connect 78 compositions in a single graph and among these, two achieve a missing connection between H6N5 and H7N6. Two alternative paths are suggested either via H6N6 or H7N5. When another CHO cells dataset is added for comparison, for example, CHO-DG44 (CVCL_7180) a cell line in which the dihydrofolate reductase (DHFR) gene locus was removed, nine new compositions are introduced while 27 are common to the first dataset. H6N6 is one of nine and the corresponding node in the new graph is transformed from virtual to full. This in turn cancels out the H*7*N5 alternative and keeps H6N6 as the only path between H6N5 and H7N6. This example illustrates how the combination of two datasets stabilises the graph connectivity.

**Figure 3:**
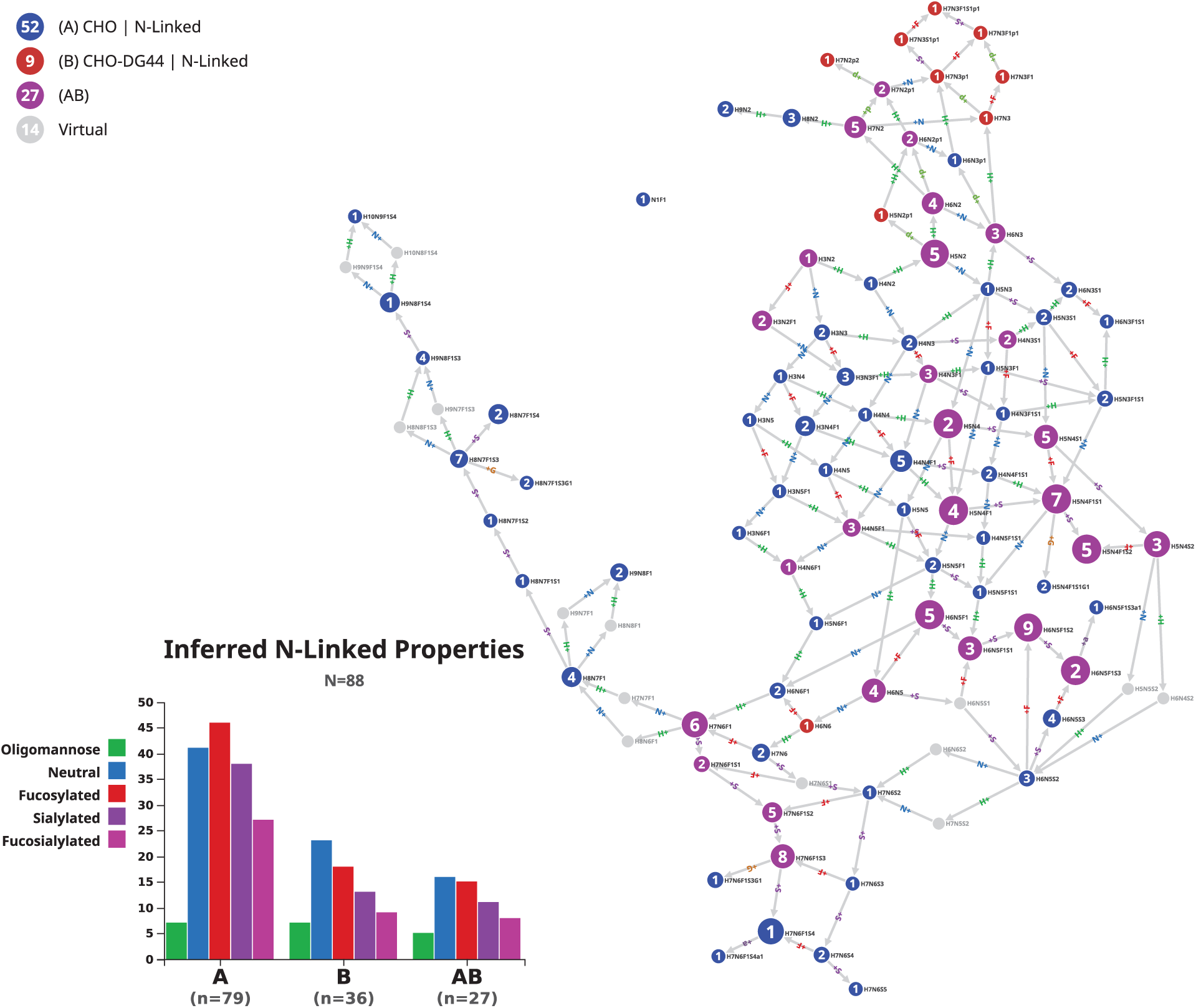
Comparison of N-glycomes of two CHO cell lines. Two N-glycome datasets of CHO cell lines are compared. Overall glycome properties are slightly different (bar plots). Common compositions are represented as magenta nodes and appear central in the graph. In relation to Figure S6 that shows graph differences arising from the introduction of virtual nodes in the N-glycome of the generic CHO cell line (CVCL_0213), the role of H6N6 is highlighted. H6N6 does not exist in CHO (CVCL_0213) but is present in CHO-DG44 (CVCL_7180) and improves connectivity.

#### N-glycome of human immunoglobulin gamma (IgG)

Automated set-ups described by several groups (18)(19)(20) have generated large datasets of the unique glycosite in peptide EEQ[F/Y]**N**ST[F/Y]R of the human IgG heavy chain (the amino acid sequence is shown here as a tryptic peptide since it appears this way in the vast majority of glycoproteomics studies). We have compiled a collection of 82 potential compositions identified on this glycopeptide (see list in supplemental material) and also used reviews such as (21)(22) that cite many relevant and useful references. Finally, additional data could be found in a recently published collaborative study (23). This dataset was input in the custom tab of Compozitor. The resulting graph is fully connected and does not require the introduction of virtual nodes.

Glycosite data described in (24) along with an HTP serum glycopeptide profiling method include results on IgG glycosylation and are stored in GlyConnect. However, this reference was not used in our compilation. The authors refer to 365 different N-glycan compositions that were entered in a customised glycan database for human serum. We did not consider this large set but only the identified intact glycopeptides of immunoglobulins reported in this article with a total of 81 compositions. This information is directly accessible in GlyConnect: https://glyconnect.expasy.org/browser/references/2857. Then, using the protein tab of Compozitor to select data corresponding to Ig gamma 1 and gamma 4, the graph generated with the 82 compositions of the compilation was enriched. Note that we did not select Ig gamma 2 because it does not add further information, nor gamma 3 that contains an additional glycosite to the regular EEQ[F/Y]**N**ST[F/Y]R.

The new graph resulting from combining these different sources is composed of 93 nodes. The 11 (=93-82) additional compositions are listed in Table 3. Interestingly, most of the corresponding nodes in the new graph are terminal or pre-terminal or they simply cause adding a single path (one arrow in, one arrow out) in the graph. In other words, they extend the graph as opposed to strengthening connectivity and as such, their removal has very limited impact on the overall topology of the graph. These nodes are circled in grey in Figure 4. Information stored in the GlyConnect corresponding records is summarised in the “comments” column of Table 3. Terminal or preterminal nodes are compositions that are either seen in other species but human, generate ambiguity with O-linked structures or were only seen in a single study. The only exception is H4N6S1. A single structure (no GlyTouCan ID available) matching H4N6S1 was described in (25) on Asn-69 of PSA (prostate specific antigen) and 32 human glycoprotein records linked to H4N6S1 originate from large-scale glycoproteomics studies. At this stage, these observations are inconclusive regarding the relevance of including this composition in the human IgG glycome.

**Table 3:**
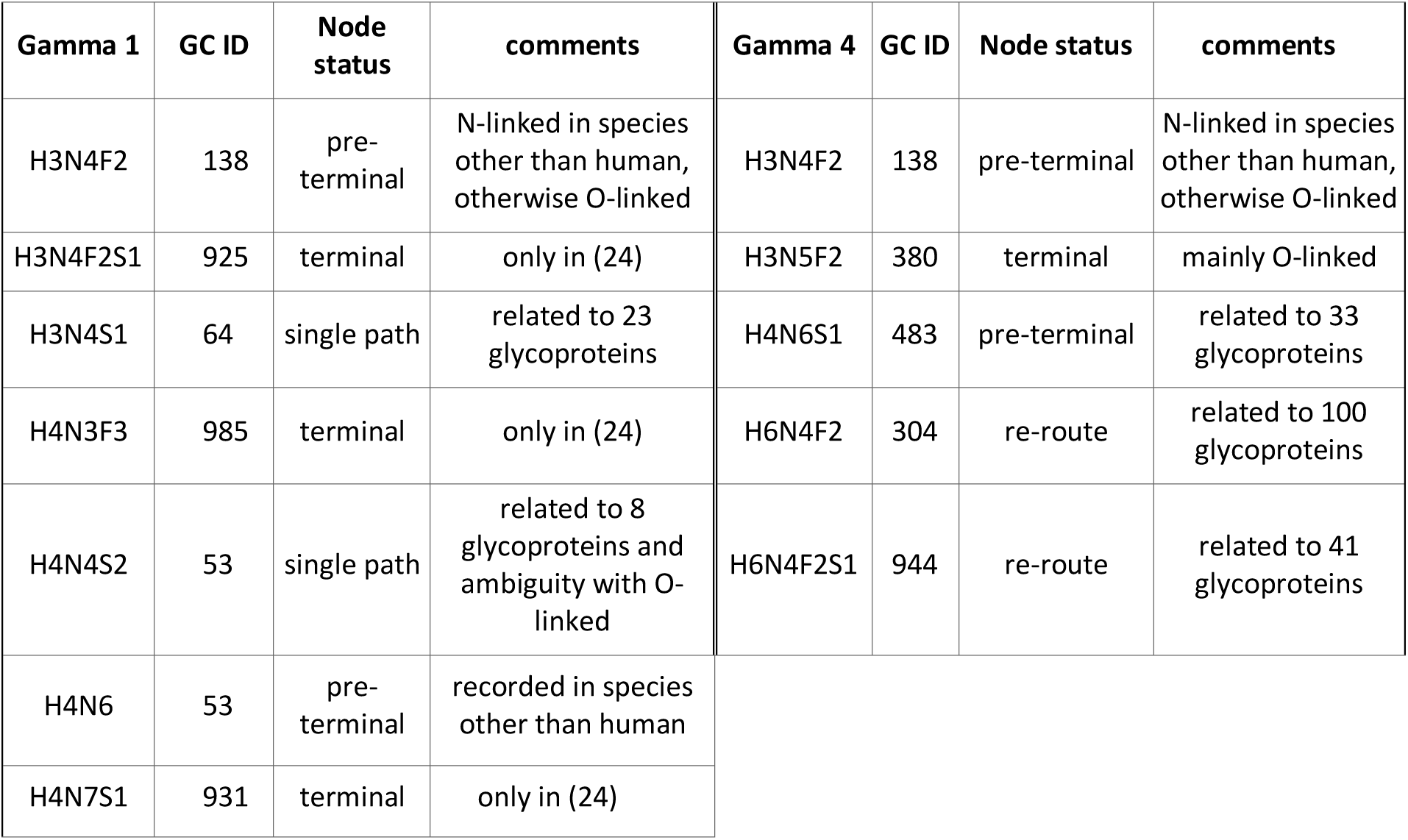
Status of new nodes in the IgG graph of combined datasets

**Figure 4:**
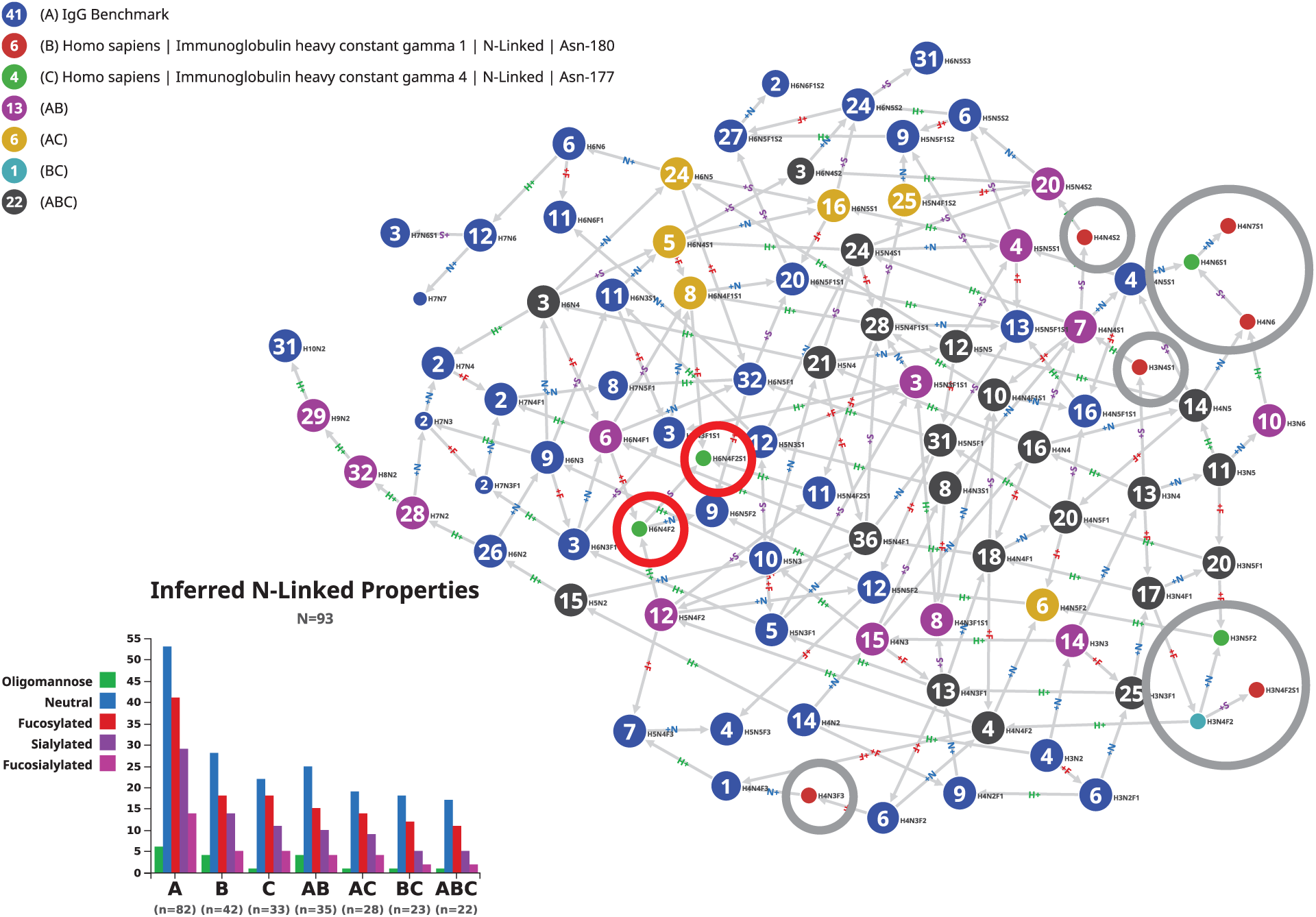
Comparison of N-glycomes of “theoretical” IgG N-glycome with two experimental datasets. Result of the combination of a custom list of 82 compositions potentially found in immunoglobulin gamma with two individual datasets submitted to Compozitor. Only ten compositions of the individual datasets are not covered by the custom list. Most of these are peripheral in the graph (circled in grey) and as such do not contribute to reinforcing connectivity. Only H6N4F2 and H6N4F2S1 (circled in red) appear as relevant connectors.

Similar uncertainty applies to the first single path node. H4N4S2 was observed as O-linked on human milk mucins and N-linked on seven human glycoproteins from large-scale glycoproteomics studies. H3N4S1 is possibly more interesting. It is reported as N-linked in human hormones from low throughput studies with two definite structures (GlyTouCan ID: G33876UV, G53933HU) and in 20 human glycoprotein records from large-scale glycoproteomics studies.

The last two additional nodes (H6N4F2 and H6N4F2S1 circled in red in Figure 4) have a stronger impact on the topology of the graph and their records in GlyConnect show more frequent occurrence. These combined features favour the inclusion of these two compositions in the human IgG glycome.

#### Tissue typing of human unspecified mucins

The GlyConnect database is populated with references that are spread over the past five decades and some earlier work was performed with limited knowledge of amino acid sequences. In that respect, the case of mucins is challenging due to the unusually large size and highly repetitive nature of these proteins particularly so, prior to the spread of high throughput DNA sequencing and the advent of genomics. As a result, a significant number of studies focused on mucosa, report glycan structures with a poor characterisation of the mucin carrier. In GlyConnect, corresponding data was harvested in 32 references published between 1980 and 2003, and are collected in a “unspecified mucin” page (GlyConnect ID: 401), which contains 251 O-linked structures matched to 102 compositions. Not a single amino acid sequence is specified, let alone a defined glycosite. Compozitor was used to visualise tissue information associated with these compositions. The “unspecified mucin” was first selected in the protein tab. Then, switching to the “source tab”, “colonic mucosa” and “pulmonary mucosa” were successively selected. The graph output of this combined query is shown in Figure 5. Both the graph and the bar plot show that most glycans of “unspecified mucins” originate from pulmonary mucosa where the domination of fucosylated glycans is obvious and in contrast with the highly sialylated colonic mucosa glycans. Admittedly, there are more compositions associated with pulmonary (82) than colonic (33) mucosa. Nonetheless, trends are observable.

**Figure 5:**
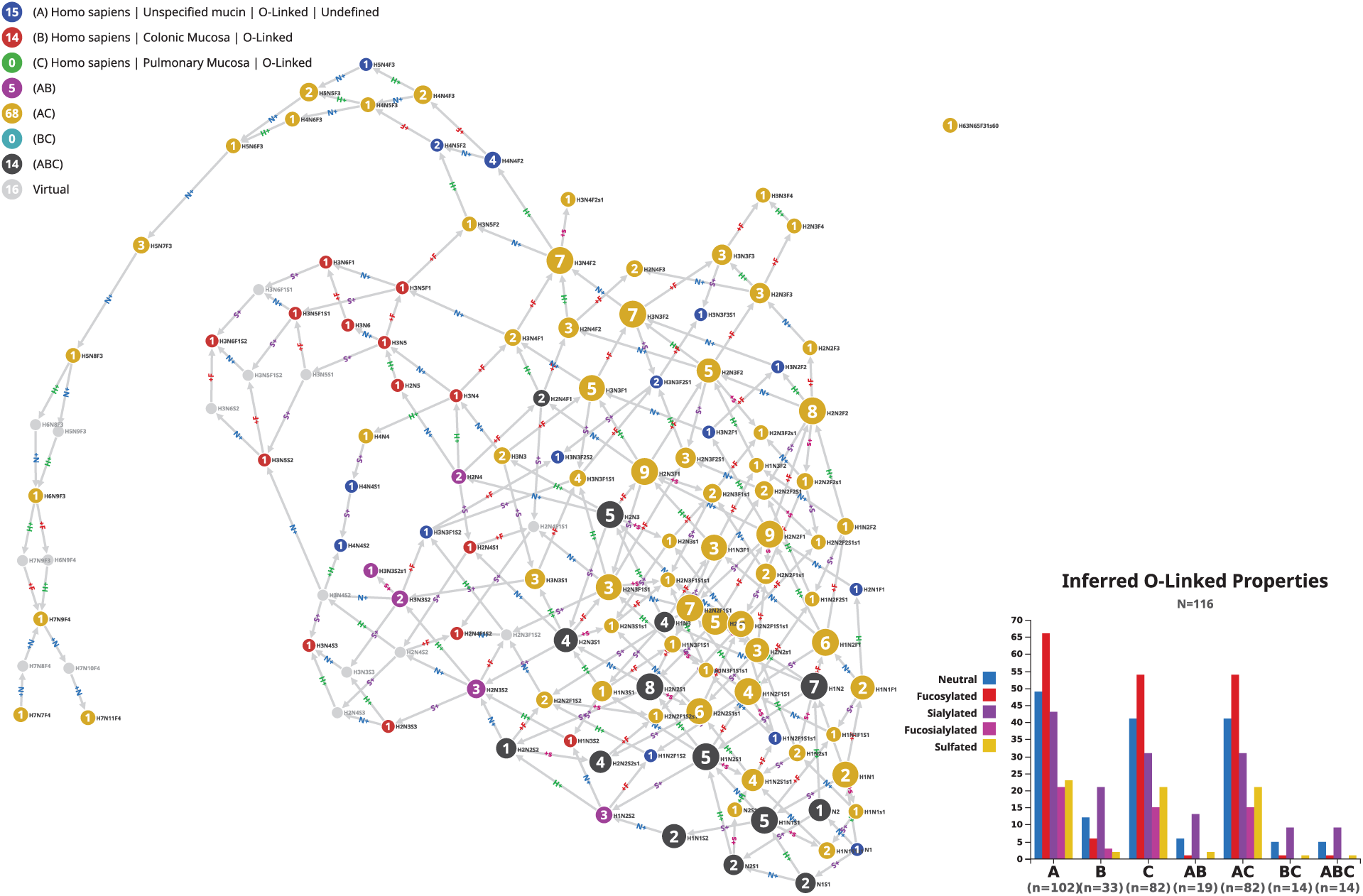
Comparison of unspecified mucin expression. Graphic result of 102 glycan compositions reported across 32 published articles involving the study of human mucosa and mucins compared with tissue information regarding mucosa as stored in the GlyConnect database. The graph shows an overwhelming presence of glycans expressed in pulmonary mucosa (yellow nodes) as opposed to colonic mucosa (red and magenta nodes). Fourteen black nodes represent compositions common to both but not highly specific of mucosa except for H2N2S2s1.

Red and magenta nodes (colonic-specific) tend to cluster together independently of yellow nodes (pulmonary-specific). Fourteen black nodes represent the compositions common to both tissues. All of these nodes have a strong outgoing connectivity except for H2N2S2s1 that is terminal and exclusive to pulmonary and colonic mucosa. Only one black node is fucosylated (H2N4F1) while the majority is sialylated. To confirm the consistency of that subset, the fourteen compositions were extracted via the Export function and pasted in the custom tab. The resulting graph (not shown) interconnects the thirteen non-fucosylated compositions with each other and only requires a virtual node from H2N3 to include H2N4F1. In the graph of Figure 5, H2N4 connects H2N3 and H2N4F1 but it appears in magenta as opposed to black, therefore it is not seen in pulmonary mucosa. It could be considered as a potential candidate to maintain consistency.

The annotation of the remaining fifteen blue nodes (non-pulmonary and non-colonic) reveals that they are spread between stomach and ovarian mucosa, milk/meconium and amniotic fluid. As summarised in Table 4, all stomach and ovarian mucosa compositions are fucosylated but not sialylated while all milk, meconium and amniotic fluid compositions are sialylated. Milk compositions are all terminal or pre-terminal. Note that N1 is the fifteenth node that is not considered in this discussion as it lacks specificity.

**Table 4:**
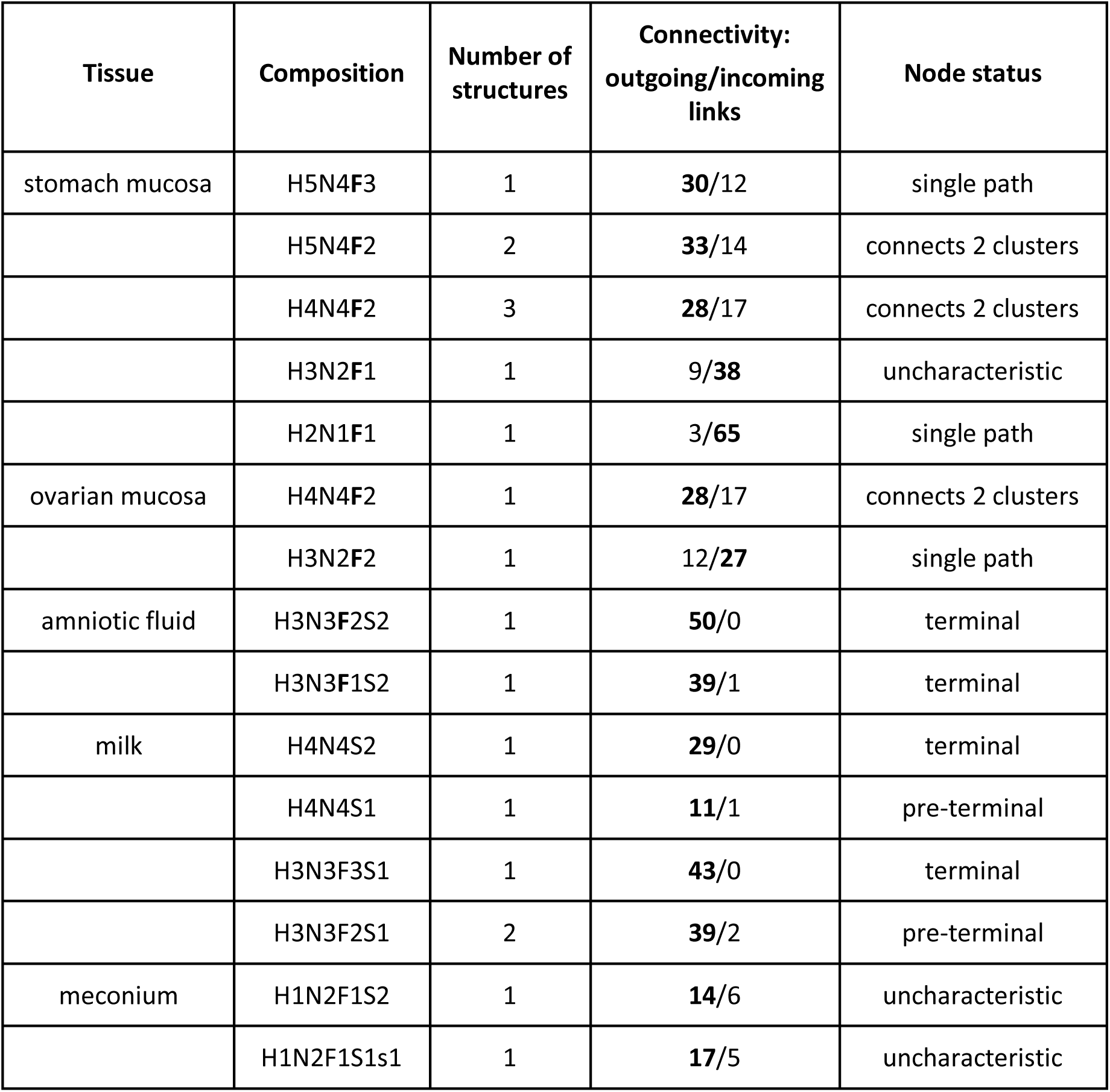
Tissue expression of mucin glycans

With this example, we show how Compozitor provides cues to refine the characterisation of a tissue glycome. It also demonstrates that data accumulation is key to identifying characteristic features. Needless to say, Compozitor graphic and interactive outputs need further interpretation and are intended as early steps in building a more refined picture of glycomes.

### Comparison of high throughput datasets

One of the key features of Compozitor is to allow for the assessment of various composition datasets that are used in intact glycopeptide identification search engines. Compozitor compares the content of these datasets with information recorded in GlyConnect so as to potentially rationalise their extension to other compositions or their reduction to rationally designed subsets. We collected three datasets: a default Byonic dataset of 305 N-glycan compositions (the default set of 309 was nowhere to be found so we reconstituted the file from supplementary material found in publications citing Byonic to reach 305 and missing four compositions), a GPQuest dataset of 181 N-glycan compositions obtained from the authors as used in (26)(27) which results are included in GlyConnect and deciphered the composition file of N-glycan compositions used with Mascot Distiller (where fucosylation is considered as a variable modification) in (3) the results of which are also integrated in GlyConnect. In the latter case, we estimated the number of possible glycan compositions to 205 given the granted possibility of accepting fucosylation as a function of the number of HexNAcs. Each of these were separately input in the custom tab of the interface to visualise the outline of the corresponding graphs and compare the respective bar plots of overall properties. These results are shown in Figure 6 along with data extracted from GlyConnect. Querying GlyConnect for all human N-glycan compositions outputs 472 items visualised in Compozitor by inputting “taxonomy=homo sapiens&glycan_type=N-linked” in the advanced tab of the interface. Note that about 60 of the additional compositions found in GlyConnect contain chemical groups such as sulphate, phosphate or acetyl that are not usually included in lists processed by search engines. Needless to say, the resulting graph is crowded and difficult to interpret yet, the summary bar plot for each case provides general information. In contrast with both Byonic and GlyConnect datasets that share a similar distribution of properties, the Mascot dataset lacks sialylated compositions along with the GPQuest dataset in which neutral compositions also seem slightly overrepresented.

**Figure 6:**
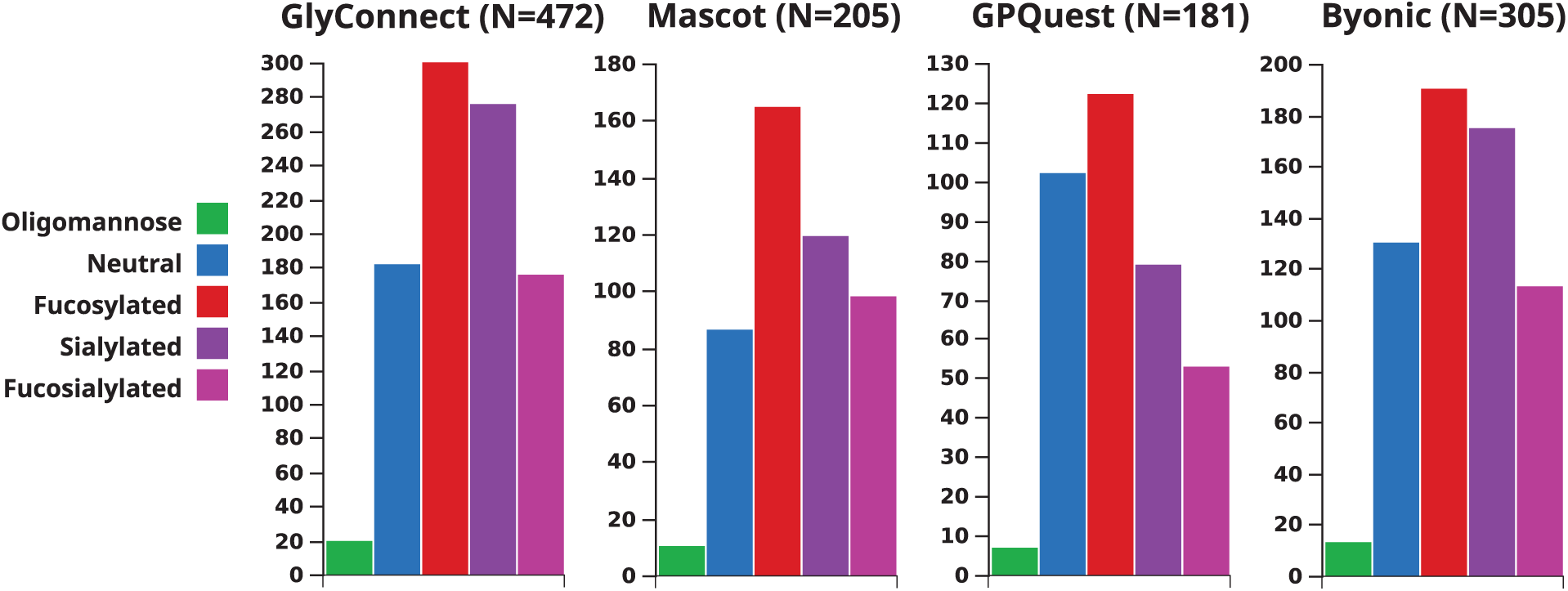
Bar plots showing properties of four selected composition files. Plotted glycan composition properties of four datasets: i) the current collection of human N-glycans in the GlyConnect database (472 in total) ii) the estimated dataset of all possible N-glycan compositions processed by Mascot Distiller in (3) (205 in total) iii) the composition file used by GPQuest in (26)(27) as communicated by authors (181 in total) and iv) the reconstituted default composition file for N-glycans in Byonic, missing 4 compositions (305 in total).

Then, to get another type of snapshot view, we compared the graph topologies of each search engine dataset as well as the effect of adding virtual nodes. To that end, we submitted each composition file to Compozitor via the custom tab. Supplemental Figures S7-S9 show the outline of the respective graphs that were generated without and with virtual nodes. The 205 compositions in the Mascot file were manifestly produced by the systematic addition of monosaccharides to the N-glycan core as seen in the regular shape and mesh-line graph (Figure S7A). As confirmed by the size and labels of the nodes, the majority of the selected structures are well documented in the GlyConnect database. Only three virtual nodes are needed to merge the two clusters created originally (Figure S7B). In contrast, many clusters characterise the GPQuest and Byonic outlines in the absence of virtual nodes (Figure S8A and S9A) and both display a strongly connected core with many side extensions when virtual nodes are included (Figure S8B and S9B). Some very large high mannose (e.g., H12N2 in GPQuest) or hybrid (e.g., H11N11S1 in Byonic) compositions remain unconnected to the main graph.

We scrutinised virtual nodes in each search engine dataset and observed that the connectivity of a specific node in different contexts is variable. Summary figures are provided in Table 5 where numbers quoted above along with the amount of compositions that are actually identified by the engines are shown. We extracted the information from GlyConnect in the original publications cited as well as two additional where GPQuest (27) and Byonic (28) were used. The ratio between theoretical (i.e., composition file provided to the search engine) and identified compositions is obviously hyper variable (from 64 to 166 in two GPQuest related articles and from 78 to 225 in two Byonic related articles) and apparently not correlated on the number of identified proteins. Further data integration is needed to appraise these variations and we are heading in this direction by planning an extensive inclusion of new data in 2020. The percentage of virtual nodes is limited and their stability may reveal relevant information on their meaning. Stability is defined here as follows: if a virtual node matches a composition already known in other contexts then this node may not be virtual in another graph and is called unstable. Conversely, if a virtual node matches a composition that never occurs in other contexts then it will remain virtual. In this latter case, the missing composition may be suggested as one to supplement a composition dataset upon the user’s judgment of its realism.

**Table 5:** Summary of composition file content in a selection of search engines

|  | Initial dataset | Identified compositions | Identified proteins | Virtual nodes |
| --- | --- | --- | --- | --- |
| **Mascot** | 205 | 77 in (3) | 257 in Ref 2 | 3 |
| **GPQuest** | 181 | 166 in (26), 64 in (27) | 934 in (26), 244 in (27) | 25 |
| **Byonic** | 305 | 225 in (4), 78 in (28) | 97 in (4), 58 in (28) | 35 |

Figure 7 shows the comparison of identified vs. possible/theoretical compositions used as input in a search engine. The version of the graph with virtual nodes was selected for that purpose. Figure 7A illustrates the case with Mascot in (3) and Figure 7B with Byonic in (4) (28). Interestingly, in Figure 7A all of the 77 compositions identified with Mascot, as confirmed by the associated bar plot, are either fucosylated or sialylated and grouped in the upper left part of the graph. Mousing over “neutral” listed in the bar plot highlights most of the blue nodes that correspond to compositions of the input file not identified in the study. In this case, the contrast between identified and unidentified is visible. In Figure 7B, 51 compositions are common to both studies using Byonic while 27 yellow nodes are specific to the endothelial cell secretome and 124 magenta nodes to prostate cancer. The bar plots indicate a steady presence of neutral compositions and a slight decrease in fucosylation. Yet, the interpretation of these results is made difficult by the gap in identification between the two studies. The remaining 53 blue nodes representing compositions not seen in any of the two cited studies are of interest if it is assumed that the reduction of the input composition file may improve the accuracy of identification. A new graph of 252 nodes was generated leaving out these 53 compositions and in which a total of 25 virtual nodes were necessary to maximise connectivity (see list in supplemental material). Five compositions were suggested as virtual nodes and matched one of the 53 “unseen” Byonic compositions: H3N6F1, H6N6, H7N7, H7N7F1S2 and H3N5F1G1. This somehow confirms their relevance in the original dataset. In contrast, twenty-three out of 53compositions contain NeuGc and could possibly be discarded when analysing human samples (note that 22/23 are fucosylated). Then, the 30 non-NeuGc containing compositions are mostly fucosylated (20/30), have either a small (<430 Da) or large (>2400 Da) molecular weight, are seldom sialylated (5/30) or contain unusual counts of the same monosaccharide (e.g., nine HexNAc or five dHex). These very specific characteristics may explain why these compositions are unmatched and deserve further attention in their selection in the first place.

**Figure 7:**
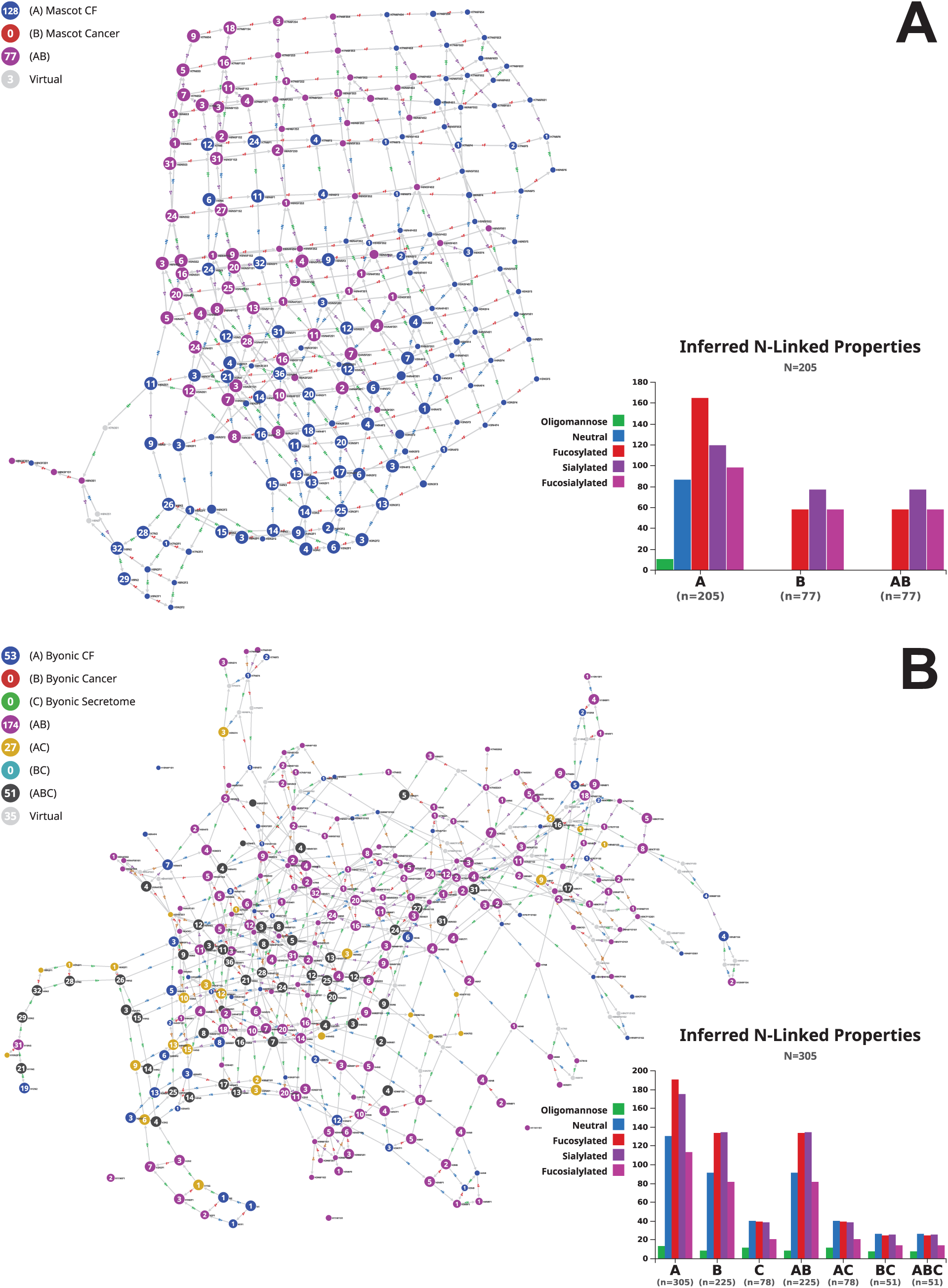
Comparison of input vs. identified N-glycan compositions in selected studies. (A) Output of the identified (magenta nodes) and estimated dataset (blue nodes) of N-glycan compositions processed by Mascot Distiller in (3) submitted to Compozitor. Magenta nodes tend to group together and cover the part of the graph where non-neutral and sialylated compositions lie. (B) Output of the N-glycan compositions identified in a prostate cancer study (4) (magenta nodes) and in the secretome of endothelial cells (28)(yellow nodes) in which the Byonic search engine was used, mapped along with our version of the Byonic default N-glycan composition file (CF) missing four items (blue nodes).

As mentioned earlier in this Results section, the connectivity and the graph location of a virtual node are criteria to be considered for determining the relevance of selecting the corresponding composition in further analyses. We selected examples in the graph of Figure 7B. H7N7 is an illustration of a potentially interesting case that questions the relevance of its inclusion when processing human data. It is not in GlyConnect yet included in the Byonic initial dataset. This means that there is no evidence of a human glycopeptide carrying H7N7 according to publications stored in GyConnect irrespective of which engine was used for identification. Nonetheless, a structure is reported in GlyTouCan (ID: G85304RG) and the composition is recorded as well (ID: G57748MK). The GlyTouCan details of G85304RG reveal that information is inherited from CarbBank (23) and two references support the existence of the structure (i) in rat kidney and (ii) in rat brain and bovine plasma. Whether to keep H7N7 in the input file remains an open question that may be context-dependent. From another angle, H5N7 is a virtual node that substantially modifies the connectivity of the graph. To illustrate this point, differential connectivity depending on the introduction of virtual nodes is highlighted in the comparative display of excerpts of the graph in Figure 7B and its counterpart without virtual nodes (full graph not shown). In Supplemental Figure S10 specific edge count (outgoing in cyan and incoming in orange) labels each displayed node. S10A (resp. S10B) is a close-up on the graph without virtual nodes (resp. with virtual nodes). The increase in edge count associated with yellow nodes (secretome data) is particularly striking with the introduction of H5N7. H5N7S1 and H5N7S2 are hardly reachable in the absence of H5N7 while the whole path from H3N2 is established if H5N7 is accounted for. Interestingly, H5N7 is recorded in the GlyConnect database (composition ID:79) but only found in chicken (29). These examples emphasise the benefit of inspecting closely a virtual node neighbourhood to determine the relevance of its inclusion.

## Conclusion

We have introduced Compozitor, a new software tool to visualise and compare glycome data based on compositional information. In the absence of a single representation of glycan compositions, the tool processes various notations and can easily accommodate new ones if necessary. Compozitor provides a range of usage through an interactive graphic interface. To begin with, it offers a different view on the content of the GlyConnect database and the option of comparing glycomes whether defined at the level of a glycosite, a glycoprotein, a cell line or a tissue. Secondly, it enables the customisation of a composition file prior to using a search engine in a glycoproteomic experiment. The content of an input file can be better rationalised from comparing glycomes. This comparison will improve with time and the planned growth of the GlyConnect database. The next major challenge is the inclusion of site-specific quantitative data. The Compozitor interface was designed to step up in that direction. Currently, the node size of a glycome graph reflects the number of associated published articles supporting the existence of a glycan composition. Once a critical mass of quantitative studies will be published, it will be easy to change this parameter and correlate the node size with the observed expression of the corresponding glycan. As other tools released as part of the Glycomics@ExPASy initiative, we also plan to improve the tool from collected feedback of users.

## Abbreviations

API: Application Programming Interface,
CHO: Chinese Hamster Ovary,
DHFR: Dihydrofolate Reductase,
ECM: Extracellular Matrix,
HUPO: Human Proteome Organization,
IgG: Immunoglobulin Gamma,
JSON: JavaScript Object Notation,
MS: Mass Spectrometry,
MS/MS: Tandem Mass Spectrometry,
REST: Representational State Transfer,
SNFG: Symbol Nomenclature for Glycans

## Acknowledgements

This work is supported by Swiss National Science Foundation [SNSF 31003A_179249]. The ExPASy portal is maintained by SIB Swiss Institute of Bioinformatics and hosted at the Vital-IT Competency Centre.

